# Behavioral Characteristics, Academic Performance, and School Enrichment for Searching Giftedness: The Case of Ambo University, Non-Boarding Special Secondary School, Ethiopia

**DOI:** 10.1101/2022.12.01.518793

**Authors:** Mengistu Debele Gerbi

## Abstract

The purpose of this study was to investigate relationship between behavioral characteristics, academic performance, and school enrichment of Ambo University non-boarding special secondary school students. To this end, correlational research was employed. Latest version of scale for rating behavioral characteristics of superior students and Purdue academic rating scale were used to generate data. 100 Ambo University Non-boarding Special Secondary School students (Male = 56, Female = 44) were selected through simple random sampling from grade 9-11. Students’ academic performances and behavioral characteristics were rated by subject teachers, homeroom teachers, and principals. Results of rating students’ academic performances revealed that students were at excellent levels in academic performance in English, mathematics, and science whereas at strong average in social studies. Results of correlation analysis showed that there was positive and significant relationship between Mathematics and English language academic performances (r = 0.332, r 2 = 0.11%, p < 0.01, df = n – 2 = 98). Mathematics performance was leading academic performance and in no subject Ambo University Non-boarding Special Secondary School students fell under average performance. 1% of students were self-nominated as gifted. Enrichment services such as health care, science technology engineering and mathematics, and instructional technologies are under satisfactory levels that need attention. Teachers and students should give emphasis to social studies as equal as other subjects.

## 1. Introduction

School enrichment is a key to enhance quality of education. A good education must balance an intended curriculum with regular, systematic enrichment opportunities that allow students to develop positive behaviors in motivation, learning, creativity, and leadership. One reason for downhill quality of education in Ethiopia might be absence of secondary schools enrichment program. Recently universities in Ethiopia have founding model secondary schools in which school enrichments are practiced. On top of this, Regional education bureaus in Ethiopia are paying attention to construct and expand model schools.

Oromia Education Bureau is dramatically expanding model schools by the name of Ifa Boru Boarding schools in many Zones of the Region in order to improve education quality. This implies that secondary schools should be enriched to heal the degenerated education quality at secondary school level. The Federal Democratic Republic of Ethiopia Ministry of Education in one hand works to achieve accessibility of education for all. In another hand the ministry strives to assure quality of education. En route practitioners of school enrichments can also search for gifted and talented students. According to the results of the current study, there are positive and significant correlation between school enrichment practices, academic performance and behavioral characteristics of outstanding. Therefore, educators, government policy-makers, educational leaders, and teachers should give due attention to secondary schools enrichment by fulfilling academic facilities such as reference books, laboratories, educational technologies, highly qualified teachers, free school meal, and conducive learning environment for all students. Educational enrichment generally for all and specifically for learners who demonstrate outstanding performance or potential for superior performance in academic, creative, leadership, and artistic domains are among popular topics all over the world. Its popularity has been increasing with an increase in societies’ awareness of the right to education for all students (15). Even though schoolwide enrichment model (SEM) originally designed as a gifted program model, currently it has been expanded as an approach used widely in schools across the world (10).

The presence of school enrichment program at non-boarding special secondary schools that supported by universities are used to protect the right to learn for outstanding students and means to realize the goal of appropriate quality education for all. Supporting high-performing students in non-boarding special school through school enrichment programs such as Science, Technology, Engineering and Mathematics (STEM) program is aimed at helping students with high academic ability to grow smartly with an astonishing base that helps them in further learning. Regarding this, (10) state that the very purpose of school enrichment is providing young people with maximum opportunities for self-fulfillment through the development and expression of one or a combination of performance areas where superior potential may be present.

Educational budget that Ethiopia has allotted for education is limited when compared with the high enrolment rate of secondary school students. Recently, about 1.5 million students are attending secondary schools in Ethiopia. It is daunting to enrich these all the secondary schools by academic facilities from the limited educational budget. As the ways out, universities in Ethiopia embarked model secondary schools by fulfilling educational facilities for high performing students. Ambo University found special non-boarding secondary school (grade 9-in 2021. The main purpose of finding such school is to assist learners in their education by closely following up their learning processes through near-teacher learning supports. Providing near-teacher learning supporting opportunities seems to provide an opportunity for academic enrichment. Accordingly, outstanding students are given the chance to attend such schools after recruited on the basis of their academic merit. A merit-based education system should be based on what one can do, rather than what one is (1).

Noticeably changed awareness of people on how important the high-achiever students are for scientific innovation, economic growth, technological advancement, and cultural progress is among factors for increasing popularity of enrichment programs (4). From the perspective of the sociology of education, special schools were once associated with elitism, but the benefits they offer for sustainable development have made them popular once again around the world.

Gifted education is among pertinent issues for countries of the world in the current global era of knowledge economy. To be competent in this global era of knowledge economy, there is concern among counties of the world with issues of academic giftedness, behavior, high academic performance of students. Terms such as excellent, superior, gifted, exceptionally able, highly able, and high-achiever are often used interchangeably in literature to refer to those students who do very well academically at school compared to their peers. The definitions of these terms overlap with one another (14). However, some studies have suggested that these terms are distinguishable and that there are differences among students categorized as gifted and high-achiever (2).

Academic giftedness is the property of being academically gifted. The definition of academic giftedness comprises of superior academic scores that manifested itself through superiority in academic learning, motivation, creativity, and leadership behaviors. According to (11), academic giftedness is schoolhouse giftedness that refers to taking tests, learning lessons, or academic giftedness. On the right-based approach, American educational law of the No Child Left Behind Act (9) states that academic giftedness is refers to students who demonstrate evidence of high achievement capability in intellectual, creative, artistic, or leadership capacity or in specific academic fields, and who need educational services that not ordinarily provided by the regular school in order to fully develop those capabilities.

The interaction between school-wide enrichment and academic performance seems to be relevant for the development of students with superior academic behavior. Research results in this area demonstrating that academically talented students who attend enrichment programs are more likely to graduate high school, attend college, and demonstrate increased knowledge and skills (12). Research on enrichment clusters documents that teachers who facilitate clusters use authentic and advanced methodologies in the clusters and then transfer those methodologies into the regular classroom teaching. An increasing body of research demonstrates increased achievement and engagement when students are exposed to challenging enrichment opportunities. However, much of the extent research has been conducted by the developers of these approaches and models, and therefore, research-based conclusions of effectiveness of enrichment on increased achievement or engagement in school, based on independent research, is generally unavailable. Hence, enriching school by providing educational facilities is necessary to develop high potentials of learners (10). The purpose of this study was to investigate relationship between school enrichment, academic performance, and behavioral characteristics of Ambo University Non-boarding Special Secondary School students.

To deal with these objectives of the current study, the following research questions were addressed. These are:

1. How teachers rate their students on academic performance behavior?

2. Is there relationship between behavioral characteristics and students’ academic performance?

3. What school enrichment program is available in Ambo University Non-boarding Special Secondary School?

## 2. Materials and Methods

### 2.1 Research Design

Mixed research approach was used for this study. In order to extract data that would most satisfactorily addresses the above research questions, researcher used quantitative research design. Both descriptive and inferential techniques of data analyses were used in this study. A correlational research method was employed to study relationship between school enrichment and students’ academic performance. Descriptive statistics were used to describe students’ behavioral characteristics. This method was used to determine association among behavioral characteristics of AUNSSS students in four subject areas (Mathematics, English, Science, and Social studies). Average academic performances for these subjects were computed to determine students’ performance characteristics. The percentage of students’ academic performance levels for each subject was also computed. Scores on scale for rating behavioral characteristics of superior students (SRBCSS) and Purdue Academic Rating Scale (PARS) were the variables considered in this study.

### 2.2 Participants

This study was conducted at Ambo University Non-boarding Special Secondary School that is located in Ambo, Western part of Ethiopia. The total population of this study was 411 (387 students, 18 teachers, 3 foreign language teachers, 1 director, 1 vice director, 1 supervisor). Simple random sampling via lottery method was used to select 100 (Male = 54, Female = 46) students. The participants of this study were selected from grade 9, 10, and 11. Twenty (20) students were selected from grade 9, forty (40) students were selected from grade 10, and forty (40) students were selected from grade 11. This was 25.8% of the total population of students. For continuous data with population size = 600, at alpha = 0.05, sample size of 100 is acceptable (8). Out of 18 teachers, 10 teachers were selected by simple random sampling. 1 director, 1 vice director, and 1 supervisor were selected by comprehensive sampling to participate in the description of school enrichment program available in the school. Sample size of the study participation was 113. The age of the participants was between 15 and 24.

### 2.3. Instruments

The instrument for academic performance characteristics was adapted from PARS that developed by (3). The items on PARS were developed by Purdue University instructors from teachers’ classroom observations, from a review of the research literature in each area, and administration of the scales that was directly derived from teachers’ classroom experiences with superior students. PARS consist of five subjects; four of them were used in the current study. The four parts of PARS used in this study were Mathematics, Science, English, and Social studies. Among five subjects on original PARS, the only set of items not included in the current study was foreign language. This exclusion was due to absence of foreign language subject other than English for grade 11students. Grade 9 and 10 students study Chinese, Arabic, and French as foreign languages. Each of the four subjects has fifteen (15) items. Thus, PARS has a total of sixty (60) items in the form of a four-scale Likert type. PARS has a well-established district line that can be expressed by a quantitative description of data for teacher-rating academic performance characteristics of students that expressed by the interval of academic performance. The possible maximum rating score for each subject is 60 whereas the minimum is 15. According to (13), the interval of academic performance or limit-line used to categorize students’ academic performance characteristics were indicated as following; below average (< 20), average (20-33), strong average (34-42), excellent (43-51), and superior (52-60). Hence, when the rating score is 25 out of 60 on a subject, student’s performance is at average. Cronbach’s alpha (*α*) reliability for the overall PARS with 60 items for the current study was *r* = 0.707. As PARS is focusing on academic performance characteristics, it is said to be a culture-fair instrument. Hence, the contextual difference would not be significant. Thus, the scale is appropriate for Ethiopian non-boarding special secondary school students.

### 2.4 Research procedures

Academic performance characteristics of students were rated by subject teachers. The teachers have got training on how to rate the students after vivid observation in the classroom and outside the class. It was assured that the teachers know each student a minimum for six months before they assigned to observe them. Based on recommendation from the manual the observation and filling the rating questionnaire took 25 days. Academic performance characteristics of science are filled by biology, physics and chemistry teachers, that of mathematics is by mathematics teachers and that of social science is by geography, history and civics teachers and that of English is by English teachers. The scores from these were summed and the average result considered.

### 2.5 Data Analysis

Raters’ responses to instruments were coded based on the 4-point Likert scale that worth Never (1), Sometimes (2), Frequently (3), and Always (4) in which 4 is the highest and 1 being the smallest rate. Using SPSS program version 23, Pearson product-moment correlation analysis was computed to determine relationships between students’ academic performance characteristics in the four subjects.

### 2.6 Ethical statement

As far as author’s responsibilities are concerned, the researcher received necessary permissions from concerned school leaders to conduct this study. Letter of a permission presented to the director of the school. After the purposes of the study were disclosed, informed permission was obtained. Investigator was also responsible to respect personal rights of participants which require protection of autonomy (privacy, confidentiality, and anonymity) and the right to know the purposes of the study. Investigator was ethically obliged not to commit any harm, and respectful to all individuals who participated in this study. Hence, any condition that suspected might have negative effects on participants was not attempted. Researcher of this study was responsible to report scientific and accurate findings of this research. Furthermore, he was responsible to duly acknowledge all authors who their materials were referred in this study and for those who their instruments have been adapted only for academic research purposes in this study.

## 3. Results

### 3.1.Results

#### 3.1.1 Descriptive Statistics of Behavioral Characteristics of Students

From Table 1, the total score on students’ learning characteristics was computed to be 5264. The mean learning characteristics of students was 52.64. Hence, 79.75% of AUNSSS students’ learning characteristics were similar to the behavior of superior students. This was calculated from number of items and the highest possible result. The number of items in this subscale is 11and the maximum possible result is 66. Then the average value of students’ learning behavior in percent is 79.75%.

**Table 1:** Learning characteristics of *AUNSSS* students as rated by 10 participants (n = 100)

| Item<br>No | Frequency of responses |  |  |  |  |  |
| --- | --- | --- | --- | --- | --- | --- |
|  | Never | Very rarely | Rarely | Occasionally | Frequently | Always |
| 1 | 0 | 0 | 7 | 32 | 42 | 19 |
| 2 | 0 | 0 | 4 | 23 | 55 | 18 |
| 3 | 0 | 0 | 3 | 30 | 52 | 15 |
| 4 | 0 | 0 | 7 | 24 | 42 | 27 |
| 5 | 0 | 1 | 6 | 27 | 42 | 24 |
| 6 | 0 | 1 | 17 | 23 | 41 | 18 |
| 7 | 0 | 0 | 7 | 33 | 40 | 20 |
| 8 | 0 | 2 | 17 | 26 | 41 | 14 |
| 9 | 0 | 0 | 5 | 27 | 37 | 31 |
| 10 | 0 | 0 | 9 | 16 | 44 | 31 |
| 11 | 0 | 0 | 13 | 15 | 47 | 25 |
| Total | 0 | 4 | 95 | 276 | 483 | 242 |
| Multiply | 1 | 2 | 3 | 4 | 5 | 6 |
| Weight | 0 | 8 | 285 | 1104 | 2415 | 1452 |
| Total score |  |  |  |  |  | 5264 |

From Table 2, the total score on students’ motivation characteristic was computed to be 5135. The mean motivational characteristic of students was 51.35. Hence, about 77.8% of AUNSSS students’ motivation characteristics were resemble to the behavior of superior students. This was calculated from number of items and the highest possible result. The number of items in this subscale is 11and the maximum possible result is 66. Then the average value of students’ motivational behavior in percent is 77.8%.

**Table 2:** Motivation characteristics of AUNSSS students as rated by 10 participants (n = 100)

| Item | Frequency of responses |  |  |  |  |  |
| --- | --- | --- | --- | --- | --- | --- |
| No | Never | Very rarely | Rarely | Occasionally | Frequently | Always |
| 1 | 0 | 5 | 10 | 23 | 44 | 18 |
| 2 | 0 | 0 | 9 | 27 | 43 | 21 |
| 3 | 0 | 0 | 9 | 27 | 49 | 15 |
| 4 | 0 | 0 | 10 | 24 | 48 | 18 |
| 5 | 0 | 0 | 9 | 36 | 41 | 14 |
| 6 | 0 | 3 | 6 | 29 | 46 | 16 |
| 7 | 0 | 0 | 12 | 30 | 40 | 18 |
| 8 | 0 | 1 | 8 | 26 | 45 | 20 |
| 9 | 0 | 3 | 11 | 20 | 38 | 28 |
| 10 | 0 | 2 | 8 | 26 | 44 | 20 |
| 11 | 0 | 5 | 15 | 29 | 36 | 15 |
| Total | 0 | 19 | 107 | 297 | 474 | 203 |
| Multiply | 1 | 2 | 3 | 4 | 5 | 6 |
| Weight | 0 | 38 | 321 | 1188 | 2370 | 1218 |
| Total score |  |  |  |  |  | 5135 |

From Table 3, the total score on students’ creativity characteristic was computed to be 4148. The mean creativity characteristic of students was 41.48. Hence, 76.8% of AUNSSS students’ creativity characteristics were look like the behavior of superior students. This was calculated from number of items and the highest possible result. The number of items in this subscale is 9 and the maximum possible result is 54. Then the average value of students’ creativity behavior in percent is 76.8%.

**Table 3:** Creativity characteristics of AUNSSS students as rated by 10 participants (n = 100)

| Item No | Frequency of responses |  |  |  |  |  |
| --- | --- | --- | --- | --- | --- | --- |
|  | Never | Very rarely | Rarely | Occasionally | Frequently | Always |
| 1 | 0 | 1 | 10 | 32 | 46 | 11 |
| 2 | 0 | 0 | 12 | 34 | 46 | 8 |
| 3 | 0 | 0 | 6 | 32 | 51 | 11 |
| 4 | 2 | 4 | 7 | 31 | 38 | 18 |
| 5 | 0 | 2 | 6 | 29 | 47 | 16 |
| 6 | 0 | 0 | 14 | 43 | 35 | 8 |
| 7 | 0 | 1 | 3 | 33 | 48 | 15 |
| 8 | 0 | 0 | 7 | 29 | 45 | 19 |
| 9 | 0 | 2 | 3 | 36 | 44 | 15 |
| Total | 2 | 10 | 68 | 299 | 400 | 121 |
| Multiply | 1 | 2 | 3 | 4 | 5 | 6 |
| Weight | 2 | 20 | 204 | 1196 | 2000 | 726 |
| Total score |  |  |  |  |  | 4148 |

From Table 4, the total score on students’ leadership characteristic was computed to be 3527. The mean leadership characteristic of students was 35.27. Hence, about 83.9% of AUNSSS students’ leadership characteristics were similar to the behavior of superior students. This was calculated from number of items and the highest possible result. The number of items in this subscale is 7 and the maximum possible result is 42. Then the average value of students’ leadership behavior in percent is 83.9%.

**Table 4:** Leadership characteristics of AUNSSS students as rated by 10 participants (n = 100)

| Item No | Frequency of responses |  |  |  |  |  |
| --- | --- | --- | --- | --- | --- | --- |
|  | Never | Very rarely | Rarely | Occasionally | Frequently | Always |
| 1 | 0 | 0 | 7 | 32 | 34 | 27 |
| 2 | 0 | 0 | 4 | 23 | 35 | 38 |
| 3 | 0 | 0 | 1 | 25 | 39 | 35 |
| 4 | 0 | 0 | 1 | 14 | 37 | 48 |
| 5 | 0 | 0 | 2 | 27 | 37 | 34 |
| 6 | 0 | 3 | 9 | 17 | 29 | 42 |
| 7 | 0 | 0 | 3 | 29 | 35 | 33 |
| Total | 0 | 3 | 27 | 167 | 246 | 257 |
| Multiply | 1 | 2 | 3 | 4 | 5 | 6 |
| Weight | 0 | 6 | 81 | 668 | 1230 | 1542 |
| Total score |  |  |  |  |  | 3527 |

Table 5 showed that the eigenvalue (E) of learning characteristics was 4.78, E of motivational characteristics = 4.66, E of creative characteristics = 4.60, E of leadership characteristics = 5.03. As indicated in Table 5 AUNSSS students exhibited about 79.8% behavior of superior students in their learning characteristics, 77.8% behavior of superior students in their motivation characteristics, 76.8% behavior of superior students in their creativity characteristics, and 83.9% behavior of superior students in their leadership characteristics.

**Table 5:** Summary of AUNSSS students’ behavioral characteristics on SRBCSS

| Domain | Score | Weighted<br>Score | Weighted<br>% value | Derived<br>S/V | Percentage |
| --- | --- | --- | --- | --- | --- |
| Learning scale | $52.64 \div 11$ | = 4.78 | X (2.5) | 11.975 | 79.75% |
| Motivation scale | $51.35 \div 11$ | = 4.66 | X (2.5) | 11.675 | 77.8% |
| Creativity scale | $41.48 \div 9$ | = 4.60 | X (2.5) | 11.5 | 76.8% |
| Leadership scale | $35.27 \div 7$ | = 5.03 | X (2.5) | 12.6 | 83.9% |
| Total |  |  |  | 47.75 |  |

#### 3.1.2 Correlation between variables of behavioral characteristics of AUNSS Students

Table 6 showed that leadership characteristics of AUNSSS students was leading behavior (M = 5.03) followed by learning characteristics (M = 4.78) and motivation characteristics (M = 4.66). Creativity characteristics was the least characteristics of AUNSSS students with (M = 4.60).

**Table 6:** Descriptive statistics for behavioral characteristics of AUNSS students (n = 100)

| Domain | Mean | Std. Deviation |
| --- | --- | --- |
| Learning characteristics | 4.78 | .66 |
| Creativity characteristics | 4.60 | .61 |
| Motivation characteristics | 4.66 | .76 |
| Leadership characteristics | 5.03 | .73 |

As Table 7 showed, there was strong positive and significant correlation between learning and creativity characteristics (r = .784, r^2^ = 61%, p < .01, df = n-4 = 96). It also showed that there was strong positive and significant correlation between learning and motivation characteristics (r = .831, r^2^ = 69 %, p < .01, df = n-4 = 96). The correlation between learning and leadership characteristics was also strong positive and significant (r = .636, r^2^ = 40%, p < .01, df = n-4 = 96). There was strong positive and significant correlation between creativity and motivation characteristics (r = .751, r^2^ = 56 %, p < .01, df = n-4 = 96). The correlation between creativity and leadership characteristics was strong positive and significant (r = .674, r^2^ = 45%, p < .01, df = n-4 = 96). It also showed that there was strong positive and significant correlation between motivation and leadership characteristics (r = .612, r^2^ = 37%, p < .01, df = n-4 = 96).

**Table 7:** Pearson product-moment correlation between variables of SRBCSS (n =100)

| S/N | Domain | 1 | 2 | 3 | 4 | Sig.<br>(2-tailed) |
| --- | --- | --- | --- | --- | --- | --- |
| 1 | Learning characteristics | 1 |  |  |  | .000 |
| 2 | Creativity characteristics | .784** | 1 |  |  | .000 |
| 3 | Motivation characteristics | .831** | .751** | 1 |  | .000 |
| 4 | Leadership characteristics | .636** | .674** | .612** | 1 | .000 |

\*\* P< 0.01 (2-tailed).

**Table 8:** Pearson product correlations between students’ academic performances (*n* =100).

| Variables | 1 | 2 | 3 | 4 |
| --- | --- | --- | --- | --- |
| Mathematics academic performance | 1 |  |  |  |
| Science academic performance | 0.298** | 1 |  |  |
| English language academic performance | 0.332** | -0.071 | 1 |  |
| Social studies academic performance | 0.158 | -0.240* | 0.194 | 1 |

\*\* . Correlation is significant at the 0.01 level (2-tailed).
\*. Correlation is significant at the 0.05 level (2-tailed).

The results of the study indicate that as academic performance in science increase, the performance in social studies decrease (r = −0.240, r^2^ = 5.76 %, p < 0.05, df = n - 2 = 98). As performance in mathematics increase, the performance in science also increase (r = 0.298, r^2^ = 8.8%, p < 0.01, df = n - 2 = 98). There is no significant correlation between Mathematics performance and Social studies performance (r = 0.158, r^2^ = 2 %, p < 0.01, df = n - 2 = 98). It also shows that there is moderate positive and significant correlation between mathematics performance and English language performance (r = 0.332, r^2^ = 11 %, p < 0.01, df = n - 2= 98).

As indicated in Table 9, 85% of AUNSSS students nominate themselves as high-achievers 14% of them as average achievers and 1% as gifted. But, the nomination by the subject teachers revealed that about 15% of the students are neither gifted nor high-achievers.

**Table 9:** AUNSSS students’ self-nomination (n = 100)

| Self-nomination category | Frequency | Percent |
| --- | --- | --- |
| High-achievers | 85 | 85% |
| Average achievers | 14 | 14 |
| Gifted | 1 | 1 |

As depicted in Table 10, teachers’ qualification in their subject area is good and interaction between students and teachers at AUNSS is at excellent level. A health care facility in the school is not available. Library facilities available in the school but there is no electronic library at the school. The school offers free school meal for students. Laboratory facilities available in the school are at good level but hands on practice method of teaching is not being exercised in the school. The school has workable learning assessment and evaluation policy to pursue learning in which the average passing mark for each subject is 70 and above. However there is a center for science, technology, engineering, and mathematics (STEM), its implementation is not functioning and at poor level. The school does not offered educational technologies such as laptop for students. Guidance and counseling services available in the school is at excellent level.

**Table 10:** Description of Enrichment Available in School as rated by school management (n= 3)

| Type of Enrichment Available in School | Level of Enrichment |  |  |
| --- | --- | --- | --- |
|  | Poor | Good | Excellent |
| Teachers' qualification in their subject area |  | √ |  |
| Level of interaction between students and teachers |  |  | √ |
| Health care facilities available in the school | √ |  |  |
| Library facilities available in the school |  | √ |  |
| Free school meal facilities available in the school |  | √ |  |
| Laboratory facilities available in the school |  | √ |  |
| Learning assessment and evaluation policy to pursue learning |  | √ |  |
| Extracurricular activities such as ESTM | √ |  |  |
| Educational technologies (e.g. laptop ) provided for students | √ |  |  |
| Guidance and counseling services available in the school |  |  | √ |

## 4. Discussion

The results of this study reveal that there are positive and significant relationships between school enrichment, behavioral characteristics and students’ academic performance.

The results from quantitative data reveal that AUNSSS students show 79% of learning behavior, 77% of motivation behavior, 76% of creativity behavior, and 83% of leadership behavior are resemble the behavior of superior students on SRBCSS. Data from SRBCSS show that AUNSSS students’ behavioral characteristics are good with 4.79, 4.67, 4.6, and 5.04 eigenvalues (Mean value on the items) for learning characteristics, motivational characteristics, creative characteristics, and leadership characteristics respectively.

The average scores for these behaviors were 52.64, 51.35, 41.48, and 35.27 respectively. Where a creative is the least, leadership is the most characteristics of AUNSS students. These results are along with the results reported by (7) in which E were 5.59, 4.27, 4.74, and 6.05 consequently for similar variables. On the other hand, the eigenivalues found in the current study are higher than the eigenvalues reported by (5) except for learning characteristics which E = 4.95. The rest eigenvalues are 2.34, 1.71, and 1.93 respectively for motivational, creative, and leadership characteristics. The current study results also show that there are strong positive and significant correlation between learning, creativity, motivational, and leadership behavior of AUNSSS students. The reason why creativity behavior is lowest behavior of the students might be due to absence of subject that focuses on creativity, hands on learning, and absence of blue-collar learning attitude.

## Conclusion

Teachers’ rating of their students on academic performance and behavioral characteristics resemble characteristics of superiors students as students at AUNSSS show 80% behavioral characteristics of superior students while 1% of students self nominated as gifted students. Positive and significant relationships are found between school enrichment, behavioral characteristics and students’ academic performance as there is increased educational enrichment in school led to increased academic performance of students. Health care facilities available in the school, extracurricular activities such as ESTM, and enrichment in educational technologies are found to poor in the school.

Based on the findings of this research, the following recommendations are suggested. AUNSSS should adapt relevant extracurricular activities in implementing the learning of students by the hands-on-learning at STEM center. Education shall be supported by 21^st^ century educational technologies. So that students shall be give laptop computers.

## Acknowledgements

The author would like to acknowledge the support provided by the affiliated school under Ambo University for providing a valuable dataset of the school to carry out this research.

## Author details

Mengistu Debele Gerbi.

PhD in Special Needs Education, Institute of education and behavioral sciences, Ambo University, Ambo City, Ethiopia.

## Data Availability

The datasets generated and analyzed in this study are available from the corresponding author on reasonable request.

## Citation information

This article could be cited as: Behavioral Characteristics, Academic Performance, and School Enrichment for Searching Giftedness: The Case of Ambo University, Non-Boarding Special Secondary School, Ethiopia, Mengistu Debele, PLOS ONE, Psychology (2023).

